# A transgenic toolkit for visualizing and perturbing microtubules in mice reveals unexpected functions for non-centrosomal arrays in epidermal morphogenesis

**DOI:** 10.1101/155994

**Authors:** Andrew Muroyama, Terry Lechler

**Affiliations:** Departments of Dermatology and Cell Biology Duke University Medical Center, Durham, NC 27710

## Abstract

Differentiation induces reorganization of microtubules (MTs) into non-centrosomal arrays in a variety of tissues. The physiological functions of these microtubule arrays are just beginning to be understood as few tools currently exist to genetically perturb microtubule organization *in vivo*, particularly in mammals. We developed a genetic toolkit that can be broadly applied to the study of microtubule dynamics and function in many cell types. Using a TRE-EB1-GFP mouse we demonstrate that distinct differentiation transitions in the epidermis cause a decrease in microtubule growth rates and microtubule growth lifetimes, resulting in strong suppression of dynamics. To understand the physiological functions of these stable, non-centrosomal microtubules, we generated a TRE-spastin mouse, which can be used to perturb microtubule organization in a wide-variety of tissues *in vivo.* Unexpectedly, microtubule perturbation exclusively in post-mitotic keratinocytes had profound consequences on epidermal morphogenesis. We uncoupled novel cell-autonomous roles for MTs in differentiation-driven cell flattening from non-cell autonomous functions in regulating proliferation, differentiation, and tissue architecture. Taken together, we have created tools that will be broadly useful for the study of microtubule dynamics and function in mammalian tissue physiology and have used them to uncover previously unknown functions for non-centrosomal microtubules during mammalian epidermal development.

## Introduction

Many studies conducted over the last several decades have provided significant insight into microtubule-associated proteins (MAPs) and their effects on microtubule dynamics and organization in cultured cells. The functions for microtubules in intact tissues, as well as their organization and dynamics, are comparatively less well-understood. There are multiple reasons for this: a) many cell types will not fully differentiate when cultured outside of the organism, b) non-specific effects resulting from using drug-induced microtubule disassembly assays, including secondary effects due to mitotic arrest and apoptosis, c) lack of information about which microtubule regulators are essential for microtubule organization in many differentiated cell types, and d) lack of genetic tools for specific perturbation of microtubules in differentiated cell types. While differentiation induces loss of centrosomal microtubule-organizing center (MTOC) activity in many cells, there are only a few examples where both the changes in microtubule organization and dynamics and the functions of the resulting networks have been assayed *in vivo* (Lacroix et al., 2014; Le Droguen et al., 2015; Oddoux et al., 2013).

The mammalian epidermis develops from a single layer of proliferative basal cells that adheres to an underlying basement membrane. As stratification proceeds, cells transit outward and progressively differentiate, first into spinous cells, then into granular cells, and finally into corneocytes, which are required for epidermal barrier activity. While microtubules are organized in a centrosomal array in basal cells, differentiation results in loss of centrosomal MTOC activity, and MTs eventually organize into cortical arrays (Lechler and Fuchs, 2007; Muroyama et al., 2016). The functions for MTs in the differentiated cells remain unknown, although they have been implicated in formation and function of cell-cell adhesions (Nekrasova et al., 2011; Simard-Bisson et al., 2017; Sumigray et al., 2012a). Here we characterize both the dynamics and the functions of epidermal microtubules using novel transgenic mouse lines that reveal differentiation-induced alterations in MT dynamics as well as unexpected functional roles for noncentrosomal MT arrays in these cells.

## Results

### Visualizing microtubule dynamics during differentiation *in vivo* with the TRE-EB1 mouse

To visualize and quantify microtubule dynamics *in vivo*, we generated a transgenic mouse containing a cassette encoding a GFP-tagged copy of the microtubule plus-tip tracking protein EB1 under the tetracycline-responsive element (TRE) promoter (Figure 1A). With this line, EB1-GFP expression can be temporally controlled through doxycycline exposure and spatially controlled via cell- or tissue-specific tTA/rtTA lines (Urlinger et al., 2000) or tissue-specific Cre lines paired with the Rosa-rtTA line (Hochedlinger et al., 2005). After screening founders by assaying EB1-GFP induction in primary keratinocytes, we chose a line with mosaic expression (hereafter referred to as TRE-EB1) to allow precise resolution of single cells in a complex tissue field.

**Figure 1:**
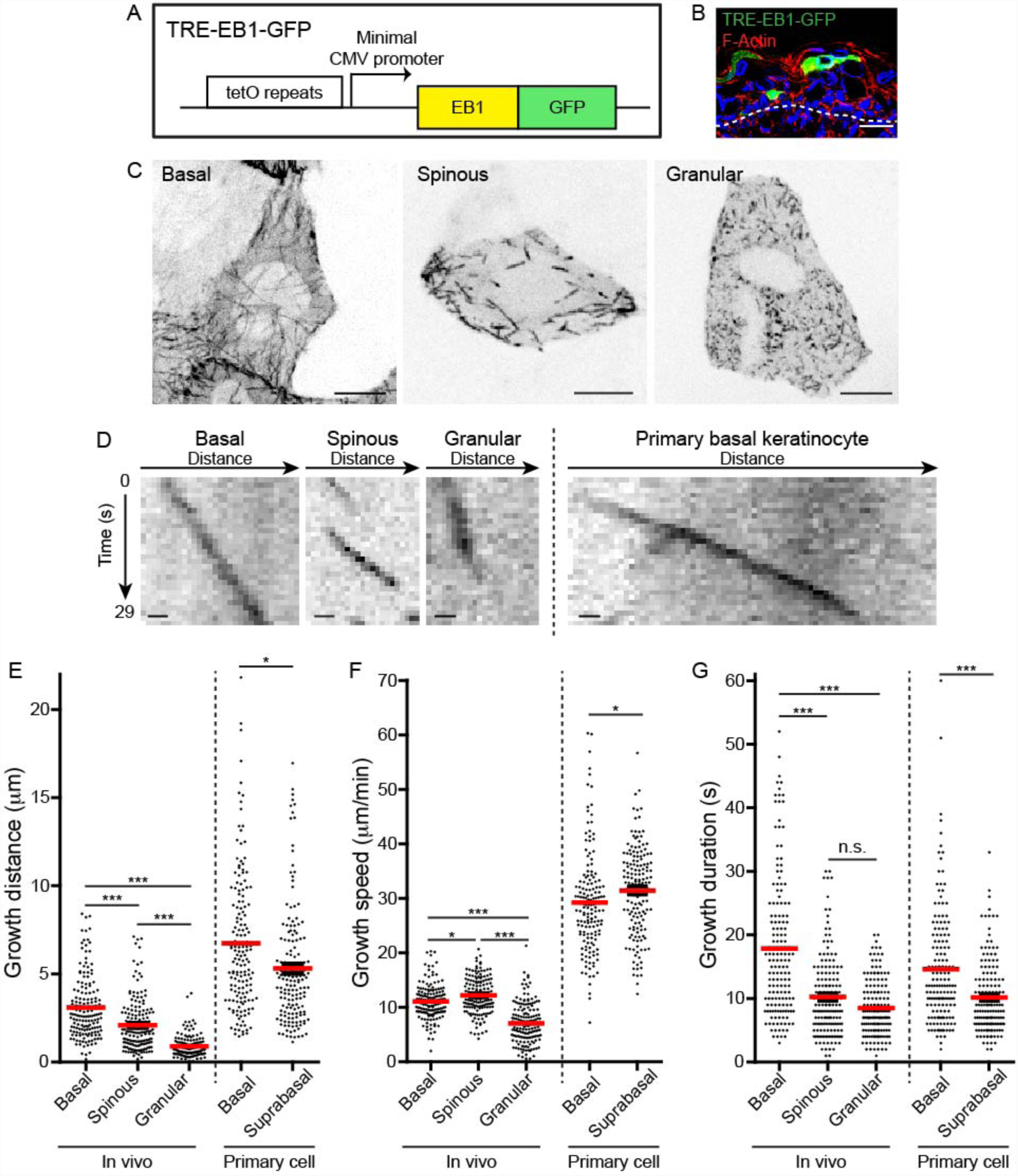
TRE-EB1 mouse line permits visualization of microtubule dynamics *in vivo.* A. Diagram of the TRE-EB1-GFP transgene. B. Cross-section of e17.5 CMV-rtTA; TRE-EB1 epidermis. Scale-20μm. C. Representative standard deviation projections of a basal, spinous, and granular keratinocyte. Scale-10μm. D. Kymographs of EB1-GFP in indicated cell types. Scale-1μm. E. Quantification of microtubule growth distance. F. Quantification of microtubule growth speed. G. Quantification of duration of microtubule growth. n=160 microtubules for each stage. Data are presented as mean±S.E.M. n.s.p>0.05, *p<0.05. ***p<0.001.

We generated CMV-rtTA; TRE-EB1 embryos to visualize microtubule dynamics in keratinocytes during progressive differentiation transitions with sub-cellular resolution (Figure 1B, Figure 1-figure supplement 1A). Importantly, we did not observe any cell morphology or tissue architecture phenotypes associated with EB1-GFP expression using this system. Microtubules in basal keratinocytes grew from the cell center to the periphery in a roughly radial orientation, in agreement with previous data that the centrosome is the primary MTOC in these cells (Figure 1C, Movie S1) (Lechler and Fuchs, 2007). In spinous cells, clear radial organization was lost, and microtubules were observed growing in all directions (Figure 1C, Movie S2). Despite the reorientation of microtubule growth in spinous cells, EB1 density was unaffected by this initial differentiation transition (Figure 1-figure supplement 1B). Similar to spinous cells, microtubules in granular cells grow throughout the cyoplasm in all directions (Figure 1C and Movie S3). However, the density of EB1-GFP puncta was higher in granular cells, and projections over time revealed that microtubules in these cells exhibited greatly reduced dynamics (Figure 1C,D). Treatment of embryos with nocodazole eliminated the GFP puncta, strongly suggesting that these EB1-GFP comets mark plus-ends of very slowly growing and/or paused microtubules (Figure 1-figure supplement 1C).

Quantification of microtubule growth distances revealed a gradual decrease in motility over keratinocyte differentiation (Figure 1E, Figure 1-figure supplement 3). Surprisingly, this gradual decrease in microtubule growth distance was due to two changes in microtubule behavior that occurred at distinct differentiation transitions. At the basal to spinous cell transition, microtubule growth speed was relatively unaffected, although there was a minor increase in growth speed in spinous cells (basal mean growth speed 11.1±3.1μm/min versus spinous mean growth speed 12.2±3.3μm/min) (Figure 1F, Figure 1-figure supplement 3). These growth rates were similar to those observed *in vivo* in cells of the *C. elegans* egg-laying apparatus and in mouse muscle (Lacroix et al., 2014; Oddoux et al., 2013). In contrast, microtubule growth speeds were strongly suppressed as spinous cells differentiated into granular cells (granular mean growth speed 7.1±3.7μm/min) (Figure 1F, Figure 1-figure supplement 3). In contrast, examination of the persistence of a single EB1-GFP puncta revealed that growth duration was significantly shorter in spinous versus basal keratinocytes (Figure 1G). The growth duration was unchanged between spinous and granular cells. The short growth periods likely reflect pause and/or catastrophe events that cannot be discriminated because EB1-GFP marks only growing microtubules. While there was no correlation between the growth speed and duration (R^2^=0.002-0.19) or the growth speed and duration (R^2^=0.004-0.2), there was a clear correlation between the length of time an EB1-GFP puncta moved and how far it traveled in basal (R^2^=0.87) and spinous cells (R^2^=0.85), as would be expected for microtubules polymerizing at a constant speed (Figure 1-figure supplement 2). Interestingly, this correlation is greatly reduced in granular cells (R^2^=0.23), (Figure 1-figure supplement 2).

To assess the importance of quantifying microtubule dynamics *in vivo*, we compared the microtubule dynamics we observed during differentiation in intact embryos to dynamics in cultured primary keratinocytes. Primary keratinocytes begin to stratify when cultured, forming a proliferative basal layer and a single, post-mitotic, “suprabasal” layer, which recapitulates some of the features of spinous cell differentiation (Muroyama et al., 2016). Absolute polymerization rates were dramatically increased in isolated cells, suggesting that microtubule dynamics are altered as primary cells initiate a wound-healing response in culture, as has been noted for cell-cell junctions (Figure 1D-G) (Foote et al., 2013). Interestingly, although the absolute polymerization rate was greatly increased, the overall trends across all three measured parameters consistently mirrored the trends seen at the basal to spinous transition *in vivo* (Figure 1E-G). These data highlight that while there is some utility in assessing microtubule dynamics in cultured cells, they do not fully recapitulate the physiological setting.

Taken together, we have established the TRE-EB1-GFP mouse line as a tool to visualize and quantify microtubule behavior in single cells *in vivo* in mice. We used this line to make qualitative observations about microtubule organization during differentiation that confirmed that microtubules reorganize into non-centrosomal arrays as basal cells differentiate. Interestingly, by following microtubule dynamics over distinct differentiation transitions, we demonstrate that pause/catastrophe events are initially increased as basal cells differentiate into spinous cells. As spinous cells mature into granular cells, microtubule polymerization rates are strongly suppressed and the occurrence of very slow growing microtubules increases significantly. To determine the functions of these distinct populations of microtubules, we next created a transgenic tool to disrupt their organization.

### Development of TRE-spastin to genetically perturb MTs *in vivo*

Because loss-of-function approaches that specifically target differentiated cells are not always feasible in tissues that rapidly turn over, we pursued a gain-of-function strategy to disrupt microtubule organization via spastin overexpression (OE). Spastin is a single-subunit microtubule severing protein whose overexpression is sufficient to dramatically perturb microtubule organization in numerous cell types (Le Droguen et al., 2015; Quintin et al., 2016; Sherwood et al., 2004). Therefore, we placed the highly active M85 spastin isoform with an N-terminal HA tag under the TRE promoter, hereafter referred to as TRE-spastin (Figure 2A) (Solowska et al., 2008).

**Figure 2:**
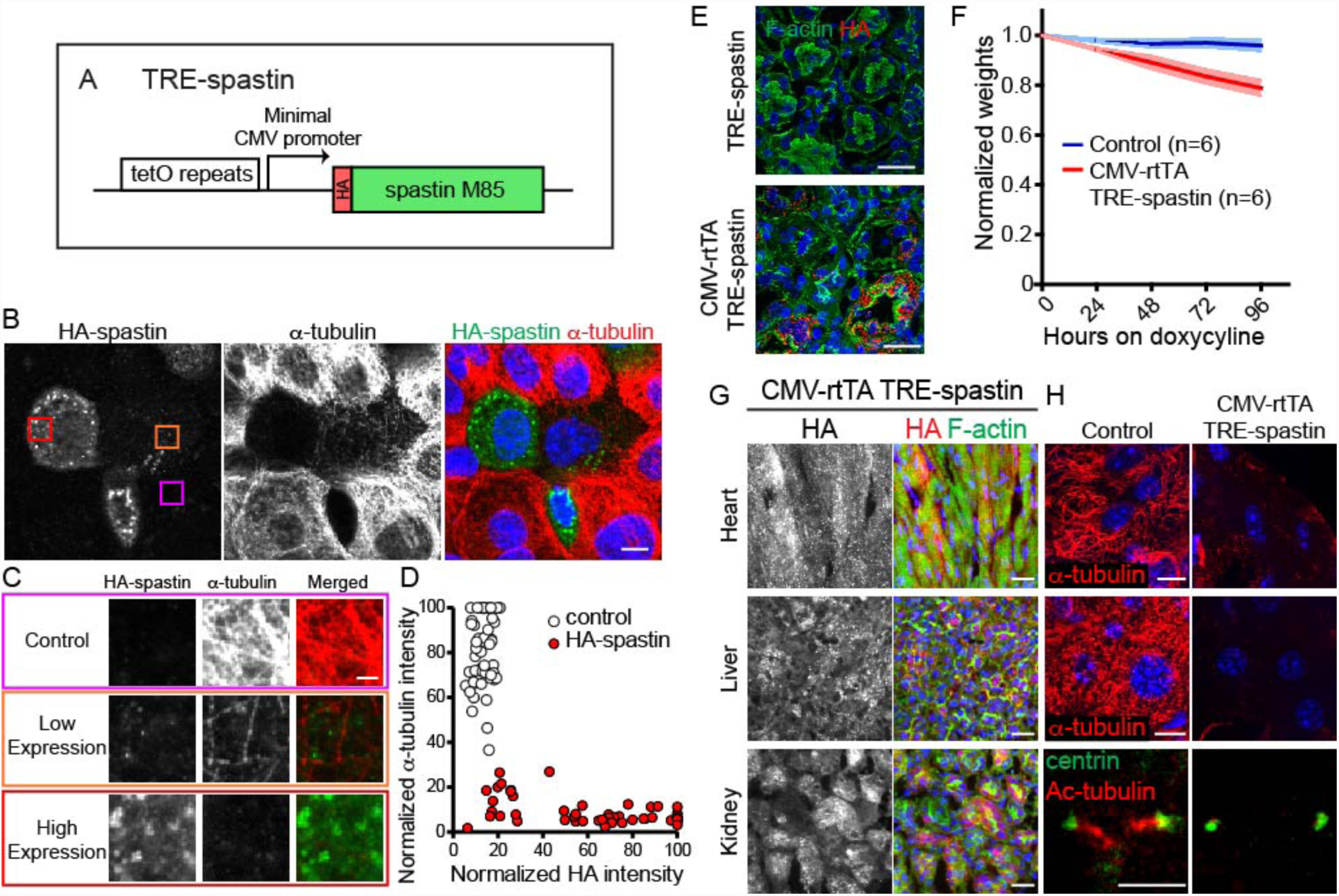
TRE-spastin expression perturbs microtubules *in vitro* and *in vivo.* A. Diagram of the TRE-spastin transgene. B. Spastin OE perturbs microtubules in cultured cells. Scale-10μm. C. Insets from (B) showing microtubule density within individual cells based on spastin expression. Scale-2μm. D. Quantification of microtubule perturbation following spastin OE. n=50 cells each from 2 independent experiments. E. Spastin expression was observed within 24 hours of doxycycline exposure and no leaky expression was detected in TRE-spastin mice. Scale-25μm. F. Weights of CMV-rtTA; TRE-spastin mice and control littermates following doxycycline exposure. G. HA-spastin expression in various tissues after 96 hours of doxycycline exposure. Scale-25μm. H. Effects of spastin OE on microtubule density *in vivo*. Note that the cilia in the kidney are dramatically shortened. Scale for the heart and liver microtubules-10μm. Scale for the cilia-5μm.

We validated the utility of this method in keratinocytes transfected with K14-rtTA and TRE-spastin plasmids. Spastin expression was only detected upon doxycycline administration, demonstrating that TRE-spastin can be temporally controlled in cultured cells. Importantly, all cells containing detectable spastin expression showed almost complete loss of microtubules, demonstrating that even low-levels of spastin OE are sufficient to severely compromise microtubule organization (Figure 2B-D).

Next, we generated TRE-spastin transgenic mice to genetically perturb MT organization *in vivo*. To validate that spastin OE can 1) be temporally controlled and 2) disrupt microtubules in tissue, we globally induced spastin OE by administering doxycycline to CMV-rtTA; TRE-spastin mice. We detected robust spastin OE in CMV-rtTA; TRE-spastin mice within 24 hours of doxycycline administration, and we did not detect HA expression in doxycycline-treated TRE-spastin mice (Figure 2E). CMV-rtTA; TRE-spastin mice rapidly lost weight and had macroscopic alterations to some organs (Figure 2F and data not shown). Importantly, spastin OE greatly reduced α-tubulin signal in all tissues assayed; often only small fragments of MTs remained following spastin induction (Figure 2G,H). In addition to cytoplasmic microtubules, spastin OE also dramatically shortened cilia (Figure 2G). Therefore, the TRE-spastin mouse can be used to genetically perturb microtubule organization *in vivo* to assess the functional roles for microtubules in numerous tissues.

### Microtubule disruption in proliferative cells of the mammalian epidermis

Next, we used the TRE-spastin mouse to understand how microtubules in distinct cell populations influence epidermal development. We generated K14-rtTA; TRE-spastin mice and induced spastin OE in proliferative basal keratinocytes (Figure 3A). Between 20-40% of basal cells overexpressed spastin using this strategy (Figure 3B). Spastin OE perturbed microtubule organization in basal cells and, as expected, induced a dramatic increase in mitotically arrested cells in mutant tissue (Figure 3C-E). Only spastin+ cells in mutant tissue had uniformly unaligned, condensed chromosomes, demonstrating that spastin OE in basal keratinocytes caused a mitotic arrest in a cell-autonomous manner (Figure 3F). Consequently, we observed increased apoptosis, presumably due to mitotic catastrophe, and a tissue-wide hyper-proliferative response to maintain progenitor number (Figure 3G,H). At this level of induction, the tissue remained architecturally normal with no detectable barrier defects, demonstrating that the epidermis is highly robust to perturbations in basal keratinocytes (Figure 3I).

**Figure 3:**
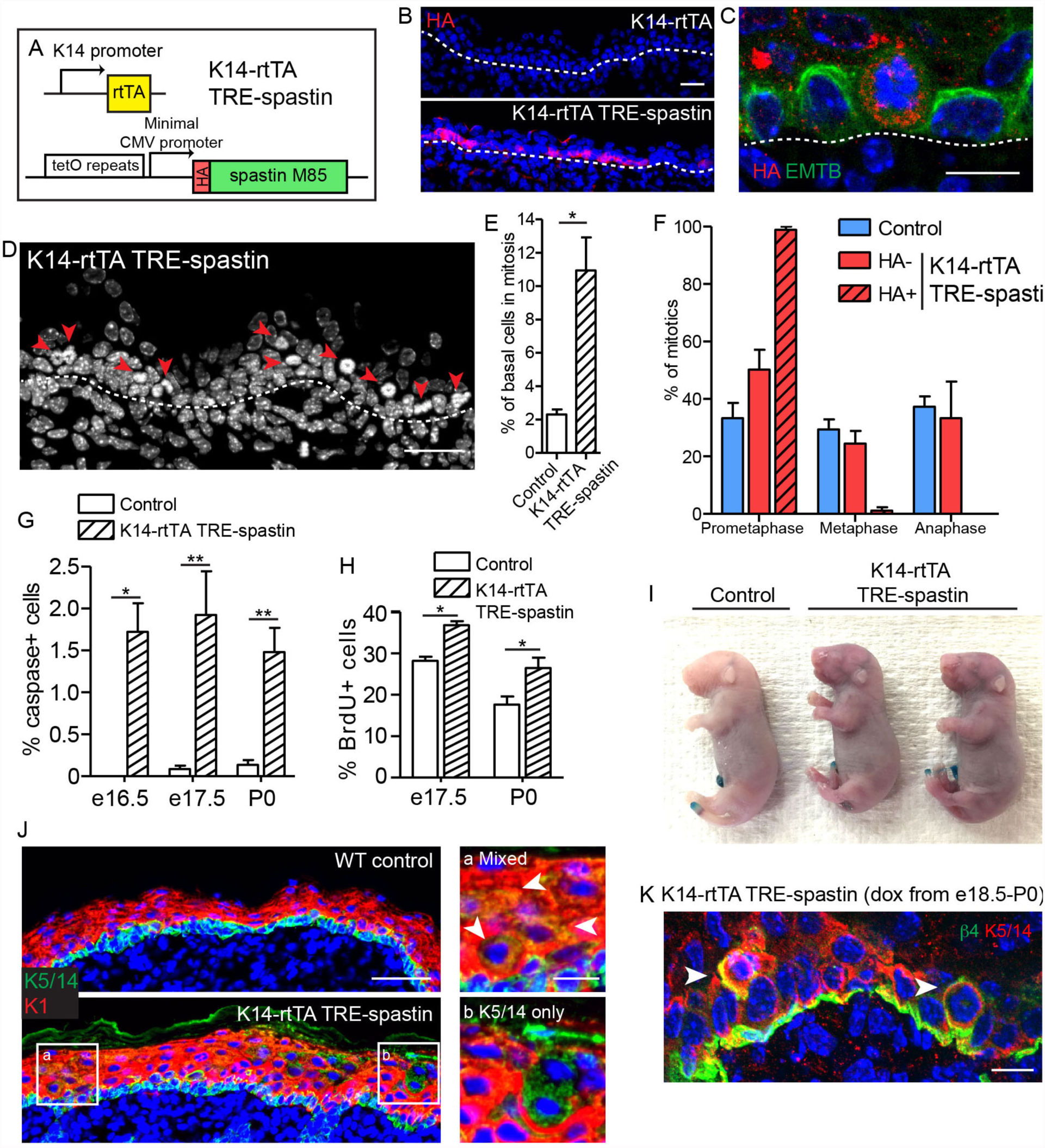
Spastin OE in basal keratinocytes induces mitotic arrest but does not alter epidermal architecture. A. Alleles used to induce spastin overexpression in basal keratinocytes. B. HA-spastin expression in e16.5 embryonic epidermis. Scale-25μm. C. Spastin OE causes microtubule loss, assayed using the 3xGFP-ensconcin microtubule-binding domain (EMTB) mouse (Lechler and Fuchs, 2007), and mitotic arrest. Scale-10μm. D. Arrows indicate mitotically arrested cells in K14-rtTA; TRE-spastin epidermis. Scale-25μm. E. Quantification of the number of basal keratinocytes in mitosis. n=3 mice per genotype. F. Quantification of mitotic stage in control back skin and spastin-negative and spastin-positive cells in K14-rtTA; TRE-spastin back skin. n=3 mice per genotype. (cell type x mitotic stage interaction, p<0.0001). G. Quantification of caspase-3-positive cells at the indicated stages. H. Quantification of BrdU+ basal cells in control and mutant back skin at the indicated stages. I. X-gal barrier assay in e18.5 embryos. J. Expression of keratins 5/14 and keratin 1 in control and K14-rtTA; TRE-spastin epidermis. Scale-50μm. Insets show zoomed regions illustrating cells expressing both K5/14 and K1 (white arrows) and also suprabasal cells that are only K5/14+. Scale-10μm. K. Delaminating mitotic cells in K14-rtTA; TRE-spastin epidermis. Scale-10μm. Data are presented as mean±S.E.M. *p<0.05. **p<0.01.

Interestingly, we noted spastin+ cells in the upper layers of the epidermis after several days of induction in the K14-rtTA; TRE-spastin line. These suprabasal spastin+ cells expressed keratin 5/14, which are normally exclusively expressed in basal keratinocytes (Figure 3J). As we have not detected any mis-expression of the K14-rtTA transgene, this result suggests that spastin+ cells delaminated from the basement membrane. Indeed, we identified many mitotically arrested cells that appeared to be in the process of delaminating, suggesting that spastin+ cells in the suprabasal layers were generated, at least partially, through delamination (Figure 3K). Thus, the epidermis eliminates mitotically arrested cells from the proliferative niche either through apoptosis or delamination. Areas with numerous suprabasal spastin+ cells displayed local thickening, suggesting that disruption of microtubules in suprabasal keratinocytes may more severely disrupt epidermal architecture (Figure 3J). Therefore, we next sought to specifically perturb microtubule organization in post-mitotic suprabasal cells, where their functions are unknown.

### Generation of K10-rtTA to specifically induce expression in suprabasal keratinocytes *in vivo*

Few tools currently exist to control transgene expression specifically in the suprabasal layers of the mammalian epidermis. Therefore, we generated a BAC transgenic in which rtTA is expressed from a large region of the keratin 10 promoter (Figure 4A). Using TRE-H2B-GFP mice, we validated that K10-rtTA faithfully recapitulates endogenous K10 expression in both embryos and adults (Figure 4). K10-rtTA induction was observed in e14.5 epidermis, as stratification commences, and was robustly and uniformly induced by e15.5 (Figure 4B) (Tumbar et al., 2004). Robust expression was observed in all K10-expressing tissues assayed, with the exception of the embryonic dorsal tongue, which only exhibited minimal induction (Figure 4C,D). Importantly, H2B-GFP expression was only detected in cells that endogenously express K10, demonstrating that our K10-rtTA line is a powerful tool to reliably control expression exclusively in post-mitotic suprabasal cells, thereby bypassing any potential mitotic abnormalities that may result from perturbation utilizing the widely adopted K14-rtTA and K5/14-CRE systems.

**Figure 4.**
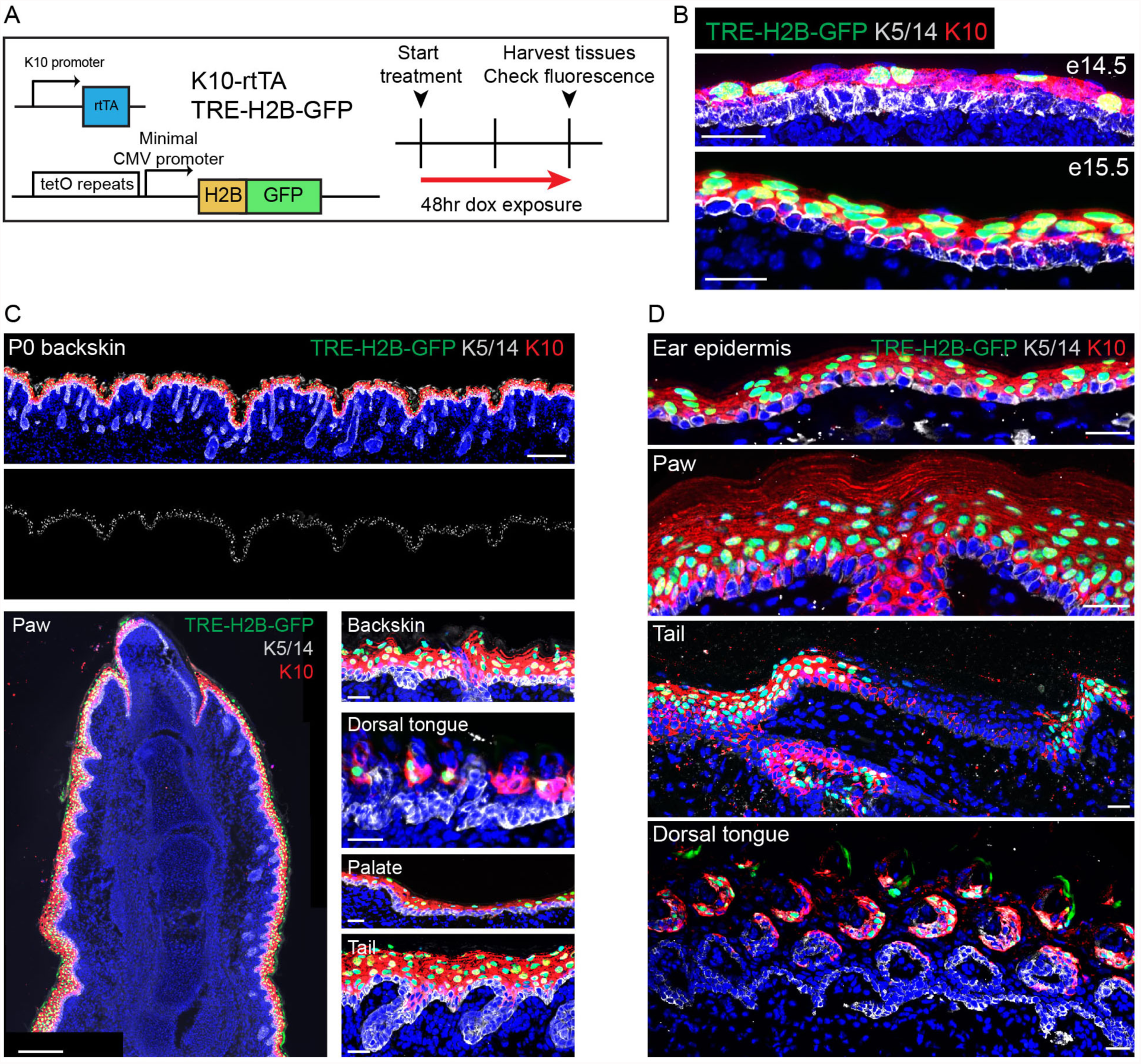
K10-rtTA expression faithfully recapitulates endogenous K10 expression. A. Alleles and experimental scheme used to validate K10-rtTA expression. B. K10-rtTA induction begins at e14.and is uniform by e15.5. Scale-25μm. C. Examples of robust K10-rtTA induction in multiple tissues in P0 pups. Scale (backskin, top)-200μm. Scale (paw, bottom left)-200μm. Scale (back skin, tongue, palate, tail, bottom right)-25μm. D. Examples of robust K10-rtTA induction across multiple tissues in adult (P30) mice. Note that in the tail, where endogenous K10 is restricted to the interscale region, H2B-GFP expression is only observed in interscale regions. Scale-25μm.

### Disruption of MTs in differentiated cells induces epidermal hyperproliferation and profound architecture defects

To understand the functions of non-centrosomal MTs in the differentiated cells of the epidermis, we used K10-rtTA; TRE-spastin mice to overexpress spastin in post-mitotic spinous and granular cells from e16.5 (Figure 5A). We confirmed that spastin expression was confined to post-mitotic suprabasal keratinocytes by HA staining (Figure 5B). Mutant neonates were recognizable from control littermates, and some exhibited flaky skin (Figure 5-figure supplement 1A). Surprisingly, spastin OE in K10-rtTA; TRE-spastin mice led to a severe thickening of the epidermis and disruptions to epidermal architecture (Figure 5C-E). Therefore, we concentrated on understanding the mechanisms by which MTs in suprabasal cells control tissue morphology.

**Figure 5.**
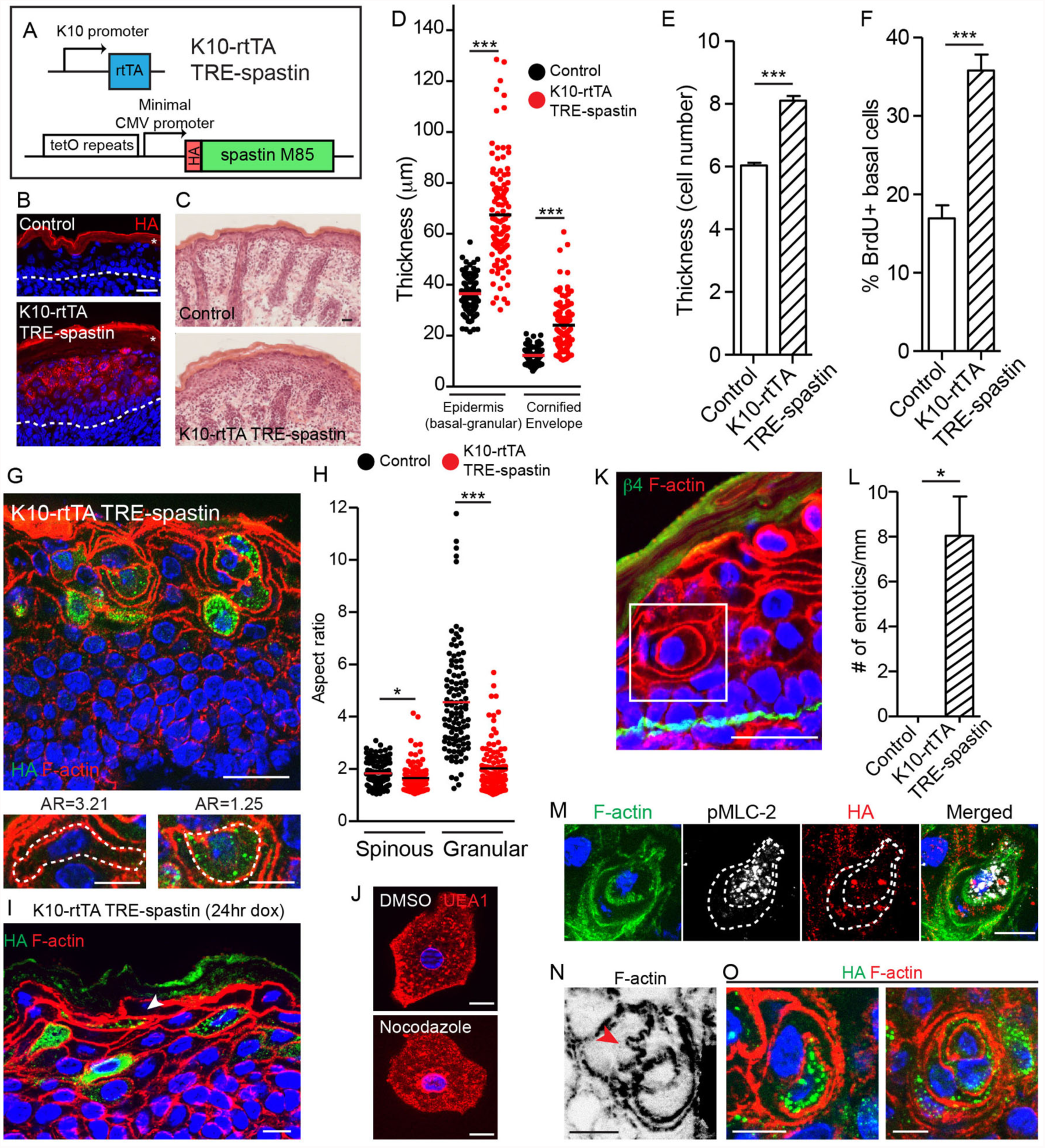
Spastin OE in differentiated keratinocytes induces cell-shape changes and entosis. A. Alleles used to overexpress spastin in suprabasal keratinocytes. B. HA-spastin expression in control and K10-rtTA; TRE-spastin epidermis. Asterisks indicate autofluorescence of the cornified envelope. Scale-25μm. C. Hematoxylin and eosin staining of control and K10-rtTA; TRE-spastin tissue. Note the cornified envelope thickness. Scale-25μm. D. Quantification of epidermal thickness in control and K10-rtTA; TRE-spastin mice. Each column is 120 measurements from 4 mice per genotype. E. Quantification of the number of cell layers present in control and K10-rtTA; TRE-spastin epidermis. n=100 measurements from 4 mice per genotype. F. Quantification of BrdU+ basal cells in control and K10-rtTA; TRE-spastin epidermis. n=4 mice per genotype. G. Cell rounding is observed in a cell-autonomous manner in K10-rtTA; TRE-spastin tissue. Scale-25μm. Zoomed regions show a spastin-negative and a spastin-positive cell within K10-rtTA; TRE-spastin tissue. Note the accompanying aspect ratios (AR). Scale-10μm. H. Quantification of the aspect ratio of individual control and spastin-positive cells. n>100 cells for each group. I. Spastin-positive granular cells after short spastin induction remain flattened. Scale-10μm. J. Isolated granular cells treated with DMSO or nocodazole. Scale-10μm. K. Example of an entotic cell in K10-rtTA; TRE-spastin epidermis. Scale-25μm. L. Quantification of the number of entotics per mm of basement membrane. n=4 mice per genotype. M. Example of an entosis where the invading cell has up-regulated phospho-myosin light chain II. The dotted line marks the cell outlines. Scale-10μm. N. Example of cell potentially invading its neighbor. Scale-10μm. O. Examples of types of entosis observed in K10-rtTA; TRE-spastin epidermis. Scale-10μm. Data are presented as mean±S.E.M. *p<0.05, ***p<0.001.

We found that the increased thickness of the mutant epidermis was associated with an increase in the number of cell layers (Figure 5E). This increase in cell number was due to a specific hyperproliferation of basal progenitor cells, as no proliferation was noted in the suprabasal cell compartment (Figure 5F). These data demonstrate a non-cell autonomous response of the progenitor cells to microtubule loss in their differentiated progeny. Hyperproliferation was not caused by elevated apoptosis, as apoptosis was not significantly increased (Figure 5-figure supplement 1B,C). Stratification occurred normally, as K5/14+ basal cells remained in a single layer above the basement membrane, but there was a dramatic thickening of the K10+ layers (Figure 5-figure supplement 1D). Taken together, our data indicate that spastin OE in differentiated keratinocytes causes dramatic hyperproliferation within the basal layer, leading to a severe thickening of all of the differentiated layers of the tissue. While barrier defects are known to induce compensatory proliferation of basal cells, we show below that the effects seen here are independent of barrier loss.

### Microtubules are required for differentiation-induced cell-shape changes

While hyperproliferation contributes to the epidermal thickening in mutant tissue, closer examination revealed that thickening was additionally driven by changes to cell shape. As keratinocytes transition from spinous to granular cells, they adopt a flattened shape that, when viewed in cross-section, is highly anisotropic (spinous mean aspect ratio (AR)=1.84; granular mean AR=4.56). Currently, little is known about how this cell-shape change is controlled. Strikingly, differentiating spastin-positive cells were incapable of properly flattening (spinous AR=1.69 versus granular AR=2.03) (Figure 5G,H). Cell-shape defects were cell-autonomous, as wild-type cells in K10-rtTA; TRE-spastin epidermis were still able to flatten, although were sometimes distorted by their spastin-positive neighbors. These cell-shape changes were not due to gross loss of adherens junctions, as we noted no obvious disruptions to cortical E-cadherin localization in mutant tissue (Figure 5-figure supplement 2). This is consistent with E-cadherin loss-of-function mutants, which are still able to undergo squamous morphogenesis (Tinkle et al., 2004; Tunggal et al., 2005). Additionally, granular cell markers were still induced in the proper layers, indicating that inability to flatten is not due to a general block in differentiation (Figure 5-figure supplement 1D). These data reveal an unexpected role for microtubules in differentiation-induced cell flattening in the epidermis. Interestingly, these phenotypes were not observed upon deletion of type II myosins or core actin regulators such as the Arp2/3 complex (Sumigray et al., 2012b; Zhou et al., 2013), suggesting that changes in granular cell morphology are not driven by alterations in acto-myosin contractility and demonstrating a role for MTs in this process.

To address whether MTs are required for initial flattening and/or maintenance of flattening, we first performed short-term treatment of K10-rtTA; TRE-spastin embryos (e18.5-P0) and measured the aspect ratio of granular cells, reasoning that many of the granular cells we observed in these back skins were flattened prior to spastin OE. Indeed, many flattened spastin+ granular cells were observed in the upper granular layers in these mice (Figure 5I). Additionally, we treated isolated granular cells with nocodazole and noted no change in cell morphology (Figure 5J). Therefore, microtubules are required for the cell flattening at the spinous to granular transition but are dispensable for maintaining the established flattened shape.

While adherens junctions appeared normal in the mutant epidermis, we were also interested in whether desmosomes, which are known upstream regulators of microtubule organization (Lechler and Fuchs, 2007; Sumigray et al., 2011), were perturbed and could underlie the cell-shape defects observed in K10-rtTA; TRE-spastin suprabasal cells. While some data in cultured cells suggests MTs are required for desmosome assembly, other studies have found little to no effect of MT disruption on desmosome formation in culture (Nekrasova et al., 2011; Pasdar et al., 1992; Simard-Bisson et al., 2017; Sumigray et al., 2011). We found a significant loss of cortical desmosomal protein staining (including Dsg1, desmoplakin, and Dsc2/3) in the spastin OE epidermis (Figure 6A). Ultrastructural analysis confirmed that desmosomes were smaller in K10-rtTA; TRE-spastin epidermis compared to control tissue (Figure 6E). Surprisingly, however, loss of cortical desmosomal components was non-cell autonomous; line-scan analysis confirmed that cortical localization of desmoplakin and Dsc2/3 was similarly disrupted between two spastin-positive, one spastin and one control, and two control cells in K10-rtTA; TREspastin epidermis (Figure 6B,C).

**Figure 6.**
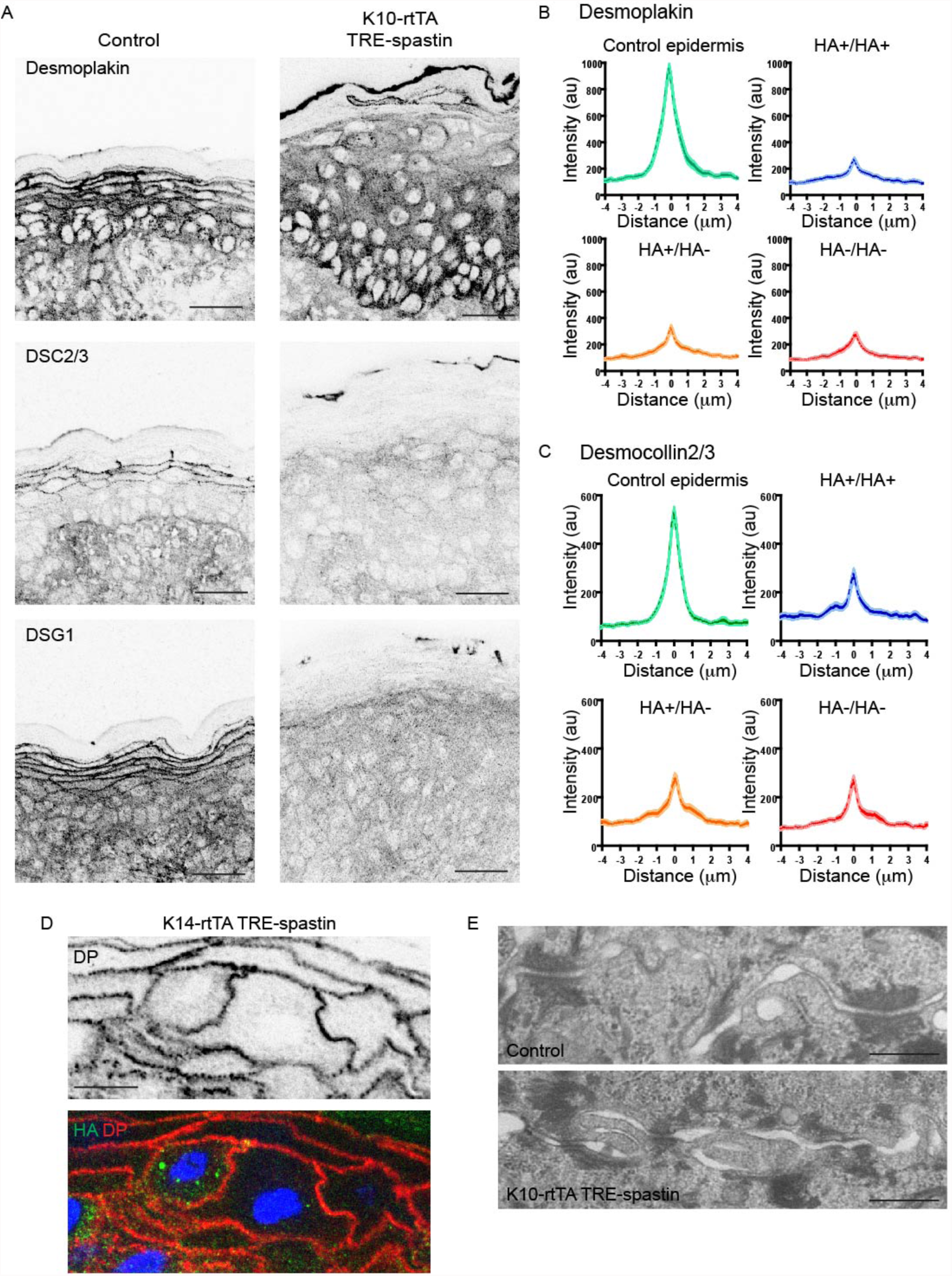
Non-cell-autonomouse desmosome defects in K10-rtTA; TRE-spastin epidermis. A. Immunofluorescence of desmosome components in control and K10-rtTA; TRE-spastin epidermis. Scale-25μm. B,C. Quantifications of desmoplakin and DSC2/3 immunofluorescence at cell-cell boundaries between indicated cell pairs. n=40 cell-cell pairs from 2 mice for each pair. D. A pair of spastin-positive cells in K14-rtTA; TRE-spastin epidermis showing that spastin expression does not intrinsically alter cortical desmoplakin localization. Scale-10μm. E. Transmission electron micrographs of desmosomes in control and K10-rtTA; TRE-spastin epidermis. Scale-500nm.

To determine whether microtubule disruption could intrinsically influence desmosome assembly, we examined pairs of spastin-positive cells surrounded by wild-type neighbors. Desmoplakin localization was normal between these single spastin OE pairs, arguing that microtubule disruption does not autonomously alter desmosome assembly in the epidermis (Figure 6D). Instead, the observed desmosome defects are not a primary effect of microtubule disruption, but rather are a tissue-wide response to it. Examination of spastin-positive cell pairs also clearly demonstrated that the cell-shape changes are not secondary to desmosome dysfunction, as these spastin OE cells with normal cortical desmoplakin levels also fail to flatten (Figure 6D). Finally, because basal cell hyperproliferation is not seen in desmoplakin-mutant skin, the progenitor hyperproliferation we observe in K10-rtTA; TRE-spastin epidermis is not likely to be secondary to desmosome disruption.

Strikingly, microtubule disruption resulted in a remarkable number of entosis-like structures in both the spinous and granular layers (Figure 5K,L). Canonical entosis is an active invasion of one cell into the cytoplasm of another as it loses substrate attachment (Overholtzer et al., 2007). We observed invading cells with elevated pMLC-2, consistent with an active actin-based entotic invasion (Figure 5M). Additionally, we observed F-actin based protrusions that appeared to be invading into neighboring cells, potentially reflecting the initial stages of entosis (Figure 5N). Entotic events ranged from entosis of a single spastin+ cell to what appeared to be concentric rings of cells, suggesting multiple layers of entosis (Figure 5O). We did not observe any examples of a wild-type cell within a spastin+ cell. Taken together, our data demonstrate that microtubules prevent entosis *in vivo*. We propose that, without microtubules, differentiated keratinocytes cannot flatten and consequently invade one another, potentially due to imbalances in membrane tension. This phenotype was not noted in desmoplakin-null epidermis, again suggesting that this is not secondary phenotype associated with adhesion defects.

The phenotypes described above – thickening of the epidermis and cell-shape defects – were also found upon disruption of microtubules in the adult epidermis, demonstrating that these phenotypes are not specific to embryonic development. Rather, microtubules are similarly essential for the homeostasis of the epidermis throughout life (Figure 5-figure supplement 3).

### Non-centrosomal microtubules are required for proper corneocyte formation but are dispensable for barrier function

Next, we wanted to assess whether the profound disruptions to epidermal architecture upon microtubule loss resulted in impaired barrier function. Epidermal barrier function is conferred through both tight junctions, which form in the granular layer, and the cornified envelopes, which are composed of enucleated, highly cross-linked corneocytes. Immunostaining for the tight junction proteins ZO-1 and occludin did not reveal any defects in spastin-positive cells or K10-rtTA; TRE-spastin tissue, demonstrating that microtubules are not required for localization of tight junction proteins in the mammalian epidermis (Figure 7-figure supplement 1A-C). Additionally, tight junctions halted biotin diffusion in mutant tissue, demonstrating that tight junction function is not observably impaired upon microtubule disruption (Figure 7-figure supplement 1D).

We noted that microtubule perturbation in K10-rtTA; TRE-spastin embryos caused formation of an abnormally thick cornified envelope (Figure 7A). By TEM, the thickened mutant CE was electron dense and many cytoplasmic remnants were observed, demonstrating a defect in corneocyte formation (Figure 7B). We confirmed that a subset of cytoplasmic proteins was retained in corneocytes in mutant back skin (Figure 7C). Retention of cytoplasmic components appeared to be cell autonomous, as we observed clear co-localization of the irregular staining with spastin+ corneocytes (Figure 7D). Isolation of the cornified envelopes confirmed that spastin OE caused severe defects in corneocyte morphology (Figure 7E,F).

**Figure 7.**
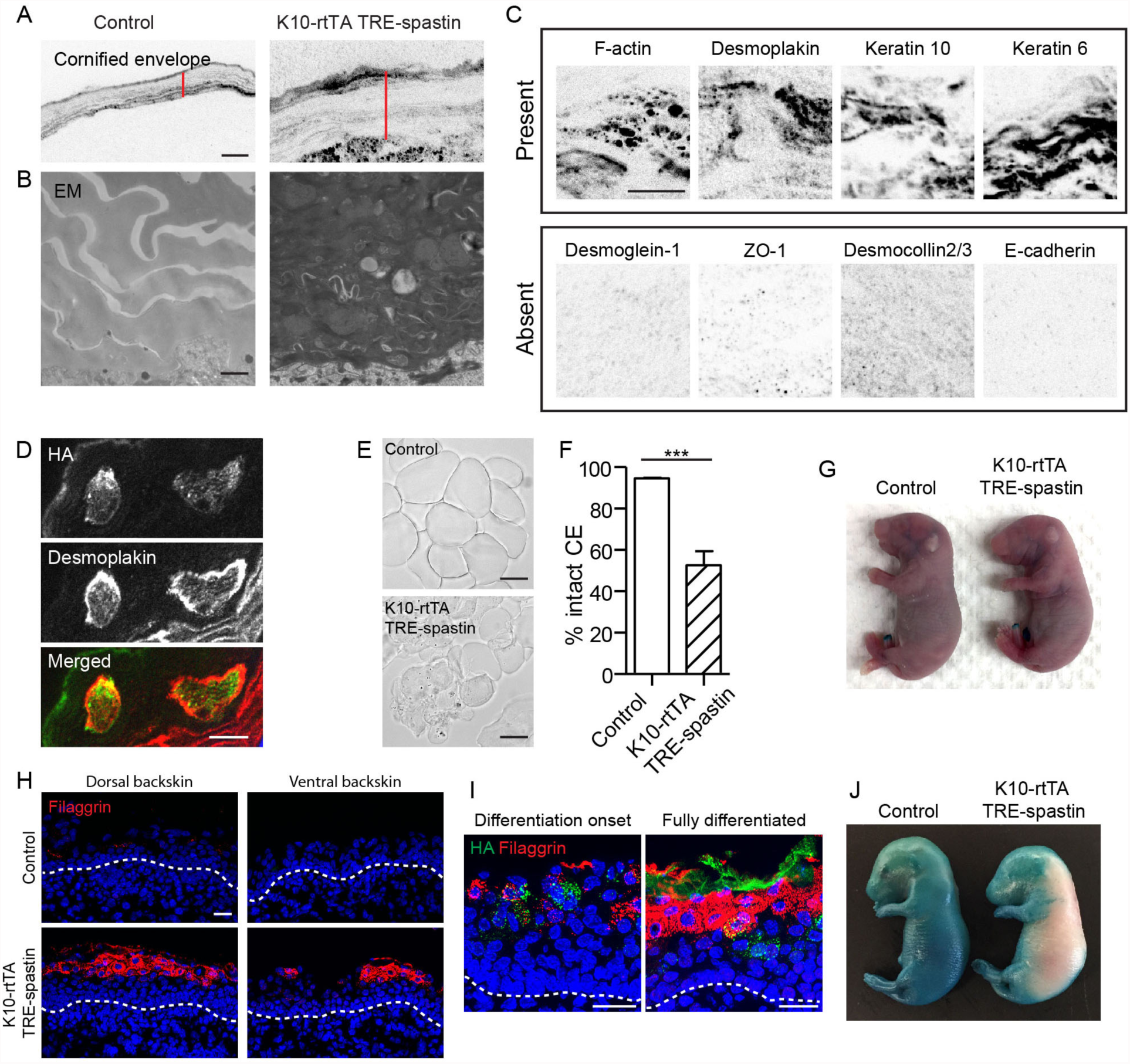
Proper corneocyte formation requires microtubules, but microtubule loss does not impair epidermal barrier function. A. CE is thickened in K10-rtTA; TRE-spastin tissue. Red lines indicate CE thickness. Scale-10μm. B. Transmission electron micrographs of cornified envelopes in control and K10-rtTA; TRE-spastin epidermis. Scale-500nm. C. Examples of protein localization in the corneocytes of K10-rtTA; TRE-spastin mice. All images are inverted fluorescence (black indicates signal). All of the indicated proteins are absent in wild-type corneocytes. Scale-10μm. D. Spastin expression cell-autonomously causes abnormal retention of cytoplasmic proteins. Scale-10μm. E. Isolated corneocytes from control and K10-rtTA; TRE-spastin mice. Scale-25μm. F. Quantification of isolated cornified envelopes. n=40 random fields from 4 mice for each genotype. G. X-gal barrier assay on e18.5 control and K10-rtTA; TRE-spastin embryos. H. Epidermal cross-sections from e16.5 control and K10-rtTA TRE-spastin embryos, stained for the differentiation marker filaggrin. Scale-25μm. I. Premature differentiation is non-cell autonomous in K10-rtTA; TRE-spastin epidermis. Filaggrin is induced in both spastin-positive and spastin-negative cells in prematurely differentiating K10-rtTA; TRE-spastin epidermis. Spastin-positive cells in the spinous layer do not induce filaggrin. Scale-25μm. J. K10-rtTA; TRE-spastin e16.5 embryos prematurely form an epidermal barrier. Data are presented as mean±S.E.M. ***p<0.001.

Because of the numerous cornified envelope defects, we performed a barrier assay to test epidermal exclusion of X-gal. Surprisingly, despite the numerous morphological defects in corneocytes, K10-rtTA; TRE-spastin embryos formed a fully functional barrier by e18.5, consistent with the fact that these mice survive postnatally (Figure 7G). We speculate that the thickening of the stratum corneum compensates for abnormal corneocyte formation. That said, our data reveal a previously unrecognized role for microtubules in the proper formation of the cornified envelope, which could either be a direct effect on corneocyte assembly or secondary to cell shape defects in granular cells.

### Microtubule disruption causes premature epidermal differentiation and barrier formation

Having identified cell-autonomous and non-autonomous functions for microtubules late in embryonic development, we turned to understand how microtubules influence the earlier stages of epidermal differentiation. We analyzed K10-rtTA; TRE-spastin embryos where spastin induction began at e14.5 as stratification commences. Surprisingly, the epidermis in mutant embryos was severely thickened even by e16.5, demonstrating that basal cell hyperproliferation is a rapid response to suprabasal microtubule disruption, even before the barrier normally forms (Figure 7H). Concurrent with thickening, mutant epidermis exhibited signs of premature differentiation, including robust expression of the granular marker filaggrin, ZO-1, and the presence of cornified envelopes (Figure 7H and data not shown). Premature differentiation was not confined to spastin+ cells; wild-type cells in K10-rtTA; TRE-spastin also expressed filaggrin in the upper layers of the tissue (Figure 7I). We performed the X-gal exclusion assay on e16.5 embryos to determine if premature differentiation in K10-rtTA; TRE-embryos resulted in premature barrier formation. Interestingly, while control embryos had no barrier function at this stage, the back skin of K10-rtTA; TRE-spastin embryos had already formed a functional barrier (Figure 7J). Therefore, microtubule disruption in suprabasal cells during early epidermal stratification unexpectedly induces premature differentiation and barrier formation, potentially through dramatic tissue thickening.

## Discussion

We generated a TRE-EB1-GFP mouse line, which permits *in vivo* imaging of microtubule dynamics in various cell types. Using this mouse, we performed live imaging of EB1-GFP in the developing epidermis of intact mouse embryos and demonstrate that microtubule growth is suppressed as keratinocytes differentiate. To our knowledge, only one other quantification of MT dynamics over differentiation in an intact tissue—the *C. elegans* egg laying apparatus—has been performed (Lacroix et al., 2014). Interestingly, specific growth parameters were tuned at distinct transitions during keratinocyte differentiation. As basal cells differentiate into spinous cells, pause/catastrophe frequency was increased, as inferred by a decrease in EB1-GFP persistence. As spinous cells matured, polymerization rates were strongly suppressed. These data raise the intriguing possibility that specific subsets of MAPs that underlie these dynamic differences may be expressed at distinct stages of keratinocyte differentiation. Mining of published transcriptional databases reveals that epidermal differentiation induces extensive alterations to the expression of microtubule-related proteins, including tubulin isoforms (α,β,γ), MAPs, and microtubule-modifying enzymes. A clear area of future work will be to determine which MAPs are responsible for the observed changes in microtubule growth parameters, with a special focus on determining how specifically tuned microtubule dynamics influence epidermal development. Furthermore, we anticipate that the TRE-EB1 mouse will be useful for performing similar measurements *in vivo* across differentiation lineages in other tissues.

We developed the TRE-spastin mouse to understand the functions for microtubules in diverse and distinct cell populations *in vivo*. This provides both the first experimental examination of microtubule function in intact epidermis and provides a microtubule “null” phenotype that is essential for future comparisons to perturbations that affect microtubule organization and/or dynamics. The dramatic consequences of microtubule disruption in the epidermis suggest that more subtle changes to dynamics/organization may have phenotypic consequences. Ablation of microtubules in different epidermal compartments revealed distinct requirements for microtubules in epidermal morphogenesis. We found that the epidermis eliminates mitotically arrested cells through both apoptosis and delamination but is quite robust in responding to microtubule defects in a substantial number of cells. The elimination of basal cells by delamination/differentiation allows the epidermis to make use of these defective progenitors.

By overexpressing spastin in suprabasal keratinocytes using a novel K10-rtTA mouse, we demonstrate that stable non-centrosomal microtubules regulate overall tissue architecture. While there are pleiotropic phenotypes downstream of microtubule loss in differentiated keratinocytes, defects in cell-shape appear to be one clear cell-autonomous consequence of microtubule disruption. In the absence of MTs, epidermal cells do not undergo squamous morphogenesis, the mechanisms of which remain poorly understood. Differentiation-induced flattening occurs in the absence of E-cadherin, type II myosins and the Arp2/3 complex, which clearly impact cell shape in numerous other cell types (Sumigray et al., 2012a; Tinkle et al., 2004; Tunggal et al., 2005). How could microtubules promote cell flattening? One possibility is that microtubules influence cell shape indirectly through other cell adhesions, such as desmosomes. However, our data strongly argue that the flattening defects we observe are independent of desmosomes, as single spastin-positive cells with normal cortical desmoplakin fail to flatten. An intriguing possibility is that kinesin-dependent microtubule sliding provides a force to drive cell-shape changes, similar to what has been reported in neurons (Winding et al., 2016). Interestingly, our data also indicate that, while microtubules are required for initiating flattening, they are dispensable for maintaining the flattened granular shape, which could be controlled by other cytoskeletal elements like keratins.

One of the clear strengths of the TRE-spastin mouse is the ability to distinguish between cell-autonomous and tissue-wide effects of microtubule disruption. Spastin OE in suprabasal keratinocytes uncovered numerous unexpected non-cell autonomous defects associated with loss of microtubules in differentiated cells. These included desmosome perturbation, basal hyperproliferation, cornified envelope defects, and premature differentiation. The mechanisms underlying these effects are a clear area for future study. Particularly, it remains important to identify how cells sense microtubule loss, whether the signal is mechanical in nature or more directly involves sensing tubulin/microtubule levels or organization. The work presented here highlights the value of examining phenotypes associated with microtubule disruption *in vivo*, and these tools should be of broad utility to the study of microtubules during mammalian development and homeostasis.

## Methods

### Mice and tissues

All mice were maintained in accordance with Duke IACUC-approved protocols. To generate the TRE-EB1-GFP transgenic mouse line, EB1-GFP was digested out of K14-EB1-GFP with SacII and NotI and ligated into pTre2 cut with the same (Muroyama et al., 2016). The XhoI site next to the SapI site in pTre2 was mutated using site-directed mutagenesis (pTre XhoI mut). Proper doxycycline-dependent expression of the TRE-EB1-GFP vector was verified in cultured keratinocytes co-transfected with a K14-rtTA plasmid and placed in doxycycline-containing media for 16 hours. TRE-EB1-GFP was linearized using XhoI and was used by the Duke Transgenic Core to generate transgenics via pronuclear injection.

To generate the TRE-spastin mouse line, the spastin M85 sequence was obtained from the pEGFP-C1 spastin M85 plasmid (kind gift of Dr. Peter Baas). First, the XhoI site within spastin M85 was mutated using site-directed mutagenesis (CTCGAG to CTAGAG) to generate a synonymous arginine mutation (amino acid 345) (CGA to AGA). An HA tag (TACCCATACGATGTTCCAGATTACGCT) was inserted at the N-terminus of the spastin M85 followed by a 4x glycine linker separating the HA tag and the start codon of spastin using PCR primers. SacII and BamHI sites were inserted on the 5’ and 3’ ends of the HA-spastin cassette, respectively. HA-spastin was PCR amplified, digested with SacII and BamHI, and inserted into pTre2 XhoI mut cut with the same. Doxycycline-dependent expression of the HA-spastin cassette was verified in cultured keratinocytes co-transfected with a K14-rtTA plasmid. The vector was linearized using XhoI and was used by the Duke Transgenic Core to generate transgenics via pronuclear injection.

To generate the K10-rtTA transgenic mouse line, the rtTA sequence was cloned behind the mouse keratin 10 promoter (BAC RP23-1D9) by the Duke Recombineering Core. Linearized DNA was used by the Duke Transgenic Core to generate transgenics via pronuclear injection.

Additional mouse lines used in this study were K14-rtTA (Nguyen et al., 2006), CMV-rtTA (Jackson labs), EMTB-GFP (Lechler and Fuchs, 2007) and TRE-H2B-GFP (Tumbar et al., 2004). For BrdU experiments, BrdU (10mg/kg) was injected into adult mice, pregnant dams (for embryos) or neonatal pups. Animals were sacrificed one hour after BrdU injection for tissue dissection and processing.

### Cornified envelope preparations

Cornified envelopes were isolated as previously described (Sumigray et al., 2011). Epidermis was isolated from P0 mice and boiled in 10mM Tris (pH 7.4), 1% β-mercaptoethanol, and 1% SDS. Remaining corneocytes were pelleted and resuspended in PBS. Resuspended corneocytes were placed on slides for imaging.

### Transmission electron microscopy

Isolated P0 back skins were fixed in EM fix buffer (2% glutaraldehyde, 4% paraformaldehyde, 1mM CaCl_2_, 0.05M cacodylate pH 7.4) for one hour at room temperature and then were placed at 4°C. Subsequent fixation, embedding, and sectioning were performed as previously described (Sumigray et al., 2011).

### X-gal barrier assay

For the X-gal barrier assay, e18.5 embryos were placed into an X-gal solution (1mg/ml X-gal, 1.3mM MgCl_2_, 100mM NaH_2_PO_4_, 3mM K_3_Fe[CN]_6_, 0.01% sodium-deoxycholate, 0.2% NP-40). After 5 hours, embryos were washed with PBS and photographed.

### Biotin barrier assay

To assess tight junction barrier function, P0 mouse pups were injected with 50ml of 10mg/ml NHS-biotin. After 30 minutes, pups were sacrificed and back skins were isolated and frozen in OCT. Biotin was detected using Streptavidin-FITC (Invitrogen).

### Cell culture

Stable wild-type keratinocytes were maintained in E low Ca^2+^ media at 37°C. Plasmid transfection was performed using the Mirus transfection reagent (Mirus).

### Granular cell isolation and drug treatment

Granular cells were isolated from P0 mice. Epidermis was isolated by placing back skin into Dispase II (2.8 units/ml) for 1hr at 37°C and 5% CO_2_. Isolated epidermis was then placed into 0.25% Trypsin-EDTA with rotating for 1hr to effectively isolate granular cells. Cells were washed with PBS, pelleted, and then resuspended in E low Ca^2+^ media with either DMSO or 10mg/ml nocodazole and placed at 37°C and 5% CO_2_ for 1hr. Cells were fixed in 4% PFA and stained with Rhodamine Ulex Europaeus Agglutinin 1 for visualization.

### Staining and antibodies

Tissue was embedded in OCT, frozen, and sectioned using a cryostat. Depending on the antibodies used, tissue sections were fixed either with room-temperature 4% PFA for seven minutes or ice-cold methanol for two minutes. Slides were washed with PBS+0.2% Triton-X and blocked with BSA, NGS, and NDS before adding the primary antibody. The following primary antibodies were used in this study: rat anti-HA (11867423001, Sigma-Aldrich), rat anti-a-tubulin (sc-53029, Santa Cruz), rabbit anti-keratin 6 (PRB-169P, Covance), chicken anti-keratin 5/14 (generated in the Lechler lab), rabbit anti-keratin 10 (905401, Covance), rabbit anti-filaggrin (905801, Biolegend), rabbit anti-loricrin (kind gift from Colin Jamera), rat anti-BrdU (ab6326, Abcam), rabbit anti-active-caspase-3 (AF835, R&D systems), rat anti-β4 integrin (553745, BD Biosciences), rat anti-keratin 8 (Troma-1, Developmental studies hybridoma bank), rat anti-ECCD2 (kind gift from Colin Jamora), rabbit anti-keratin 1 (kind gift from Colin Jamora), mouse anti-desmoplakin (CBL173, Chemicon/Millipore), mouse anti-desmocollin-2/3 (clone 7G6, Santa Cruz), mouse anti-desmoglein-1 (610273, BD Biosciences), rabbit anti-phospho-myosin light chain 2 (Thr18/Ser19) (3674, Cell Signaling), rabbit anti-occludin (ab3172, Abcam), rabbit anti-ZO-1 (61-7300, Zymed/Invitrogen), rabbit anti-centrin1 (ab101332, Abcam), mouse anti-acetylated tubulin (T7451, Sigma-Aldrich), Rhodamine Ulex Europaeus Agglutinin 1 (RL-1062, Vector Laboratories). F-actin was visualized using fluorescently conjugated Phalloidin (A12379, Invitrogen and P1951, Sigma-Aldrich).

### Image acquisition

Images of the K14-rtTA; TRE-spastin keratinocytes and mice were acquired on a Zeiss Axio Imager microscope with Apotome attachment with the following objective lenses: 20x Plan-Apo 0.8 NA lens, 40x Plan-Neofluar 1.3 NA oil lens, and 63x Plan-Apo 1.4 NA oil lens. Images were acquired using AxioVision software. All images of the K10-rtTA; TRE-spastin mice were acquired on a Zeiss Axio Imager microscope with Apotome 2 attachment and Axiocam 506 mono camera with the same objectives. When making intensity measurement comparisons, all images within one experiment were taken with identical exposure times.

Movies of EB1-GFP in CMV-rtTA; TRE-EB1 embryos were acquired on an Andor XD revolution spinning disc confocal microscope at 37°C and 5% CO_2_ using a 60x Plan-Apo 1.2 NA water objective. Briefly, embryos (e11.5 (for basal), e15.5 (for spinous), or e17.5 (for granular)) were dissected and placed onto glass-bottom dishes (MatTek) in E media for imaging. EB1-GFP movies were acquired with at 1 frame per second. Embryos were used for a maximum of one hour after isolation. Images on the spinning disc microscope were acquired using MetaMorph software. For imaging EB1-GFP in isolated, primary keratinocytes, epidermis was isolated from P0 back skin by overnight incubation in dispase (1U/ml). Keratinocytes were isolated by incubation in 1:1 0.25% trypsin-EDTA:versene and plated onto glass-bottom dishes in E no Ca^2+^ media supplemented with 0.5mM Ca^2+^ for 48 hours before imaging. Images of EB1-GFP in isolated, primary keratinocytes were acquired on a Leica DMI6000 microscope at 37°C and 5% CO_2_ using a 63x Plan-Apo 1.4-0.6 NA oil objective. These images were acquired using SimplePCI software. For DMSO and nocodazole experiments, embryos were first screened for EB1-GFP expression on the Leica DMI6000 microscope and then placed into E media with 10mg/ml nocodazole or DMSO for 30 minutes at 37°C and then imaged on the spinning disc microscope.

### Image quantification

All image quantification was done using FIJI software. EB1-GFP comets were tracked manually by marking the starting and ending positions of an EB1 comet and noting the number of frames during which the comet was visible. All EB1 tracks were considered to be straight lines, and any that deviated sharply or curved were excluded from the analysis. For quantifying EB1 comet density, the number of EB1 comets was counted and the cell cytoplasm was manually outlined to calculate the cellular area. For quantification of α-tubulin intensity in K14-rtTA; TRE-spastin keratinocytes, each cell was outlined and the average HA and α-tubulin intensities were obtained for each cell. The mean intensities for each cell were then normalized to the maximum value in the picture (HA from spastin+ cells and α-tubulin from control cells). Aspect ratios for the differentiation-induced cell shape changes were calculated by tracing individual cells and using the measurement option in FIJI. Line-scan analysis of desmoplakin and desmocollin2/3 was performed by manually drawing 5 pixel-wide lines across cell-cell boundaries and calculating the mean fluorescence intensities at each point along the lines. Maxima were aligned and the ends were trimmed to yield the final line scan. All statistical analysis was performed using GraphPad Prism 5 software.

## Acknowledgments

We thank Peter Baas for reagents, Julie Underwood for care of the mice, and members of the Lechler Lab for valuable advice. In addition, we thank the Duke Transgenic Core for the generation of novel mouse lines. This work was supported by an NSF predoctoral grant to AM and by grants from the NIH (NIAMS and NIGMS), R01AR055926, R01AR067203 and R01GM111336 to TL.

## Author Contributions

A.M. performed the experiments. A.M. and T.L. designed the experiments and wrote the manuscript.

---

**Figure 1-figure supplement 1.**
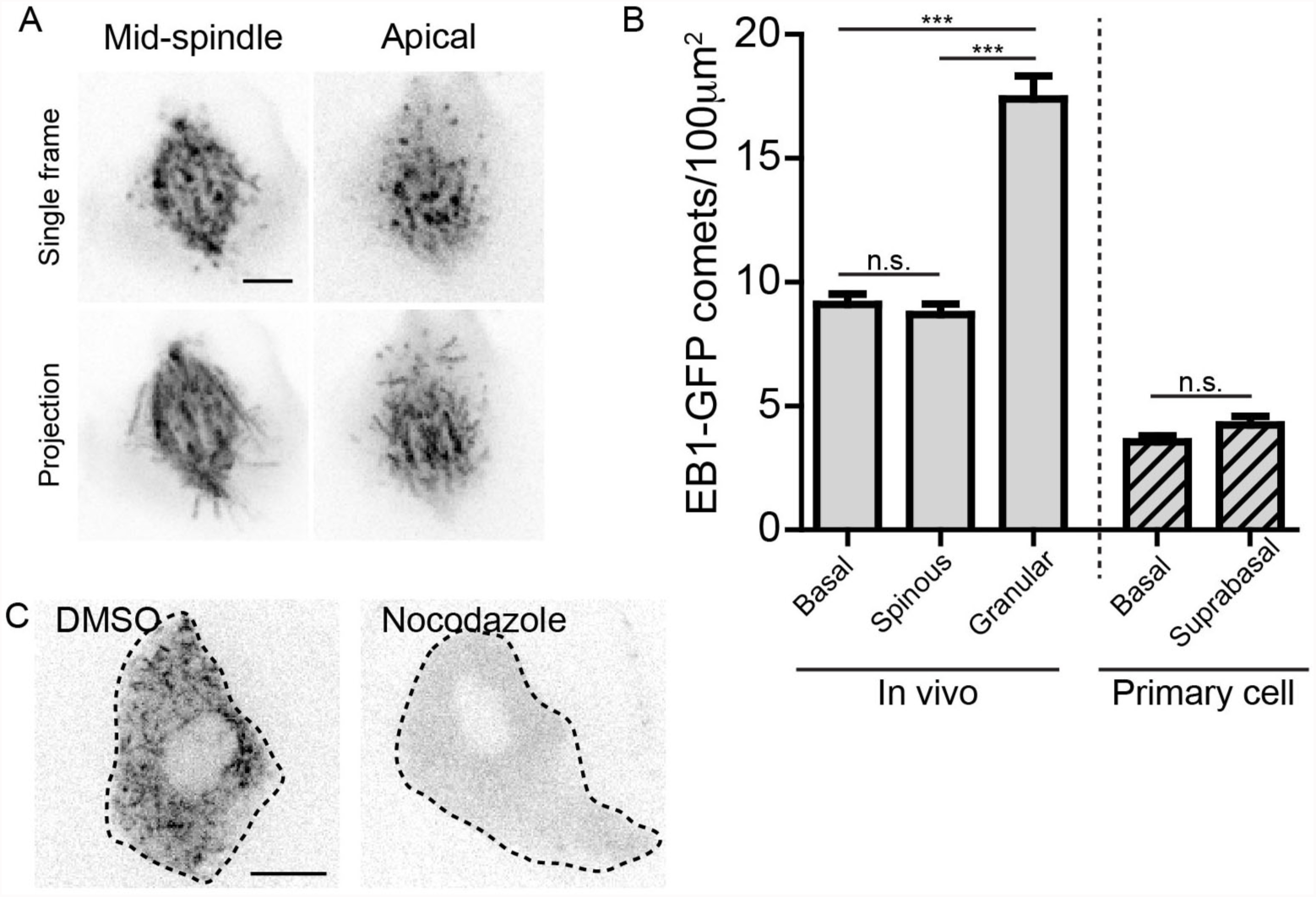
Validation of EB1-GFP line and characterization of microtubule density. A. EB1-GFP labeling a mitotic spindle at sub-cellular resolution in a basal keratinocyte. Scale-5μm. B. Quantification of EB1-GFP density in indicated cell types. n=25 cells for each cell type. C. Single frame of EB1-GFP in granular cells in e17.5 embryos treated with either DMSO or nocodazole. Scale-10μm. Data are represented as mean±S.E.M. n.s.-p>0.05, ***p<0.001

**Figure 1-figure supplement 2.**
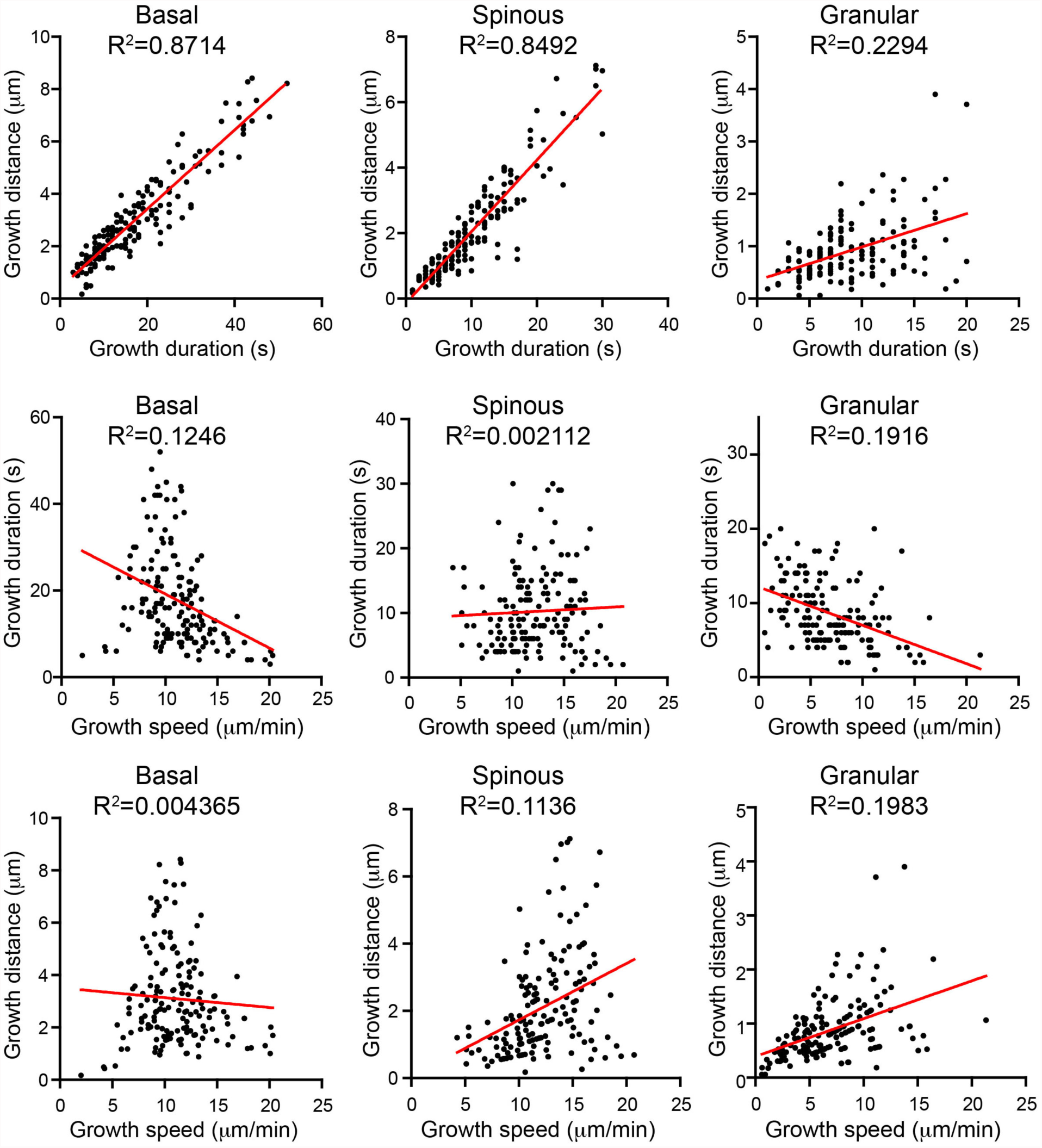
Correlation analysis of microtubule parameters for individual microtubules in the indicated cell types. Note the loss of correlation between growth distance and duration in granular cells.

**Figure 1-figure supplement 3.**
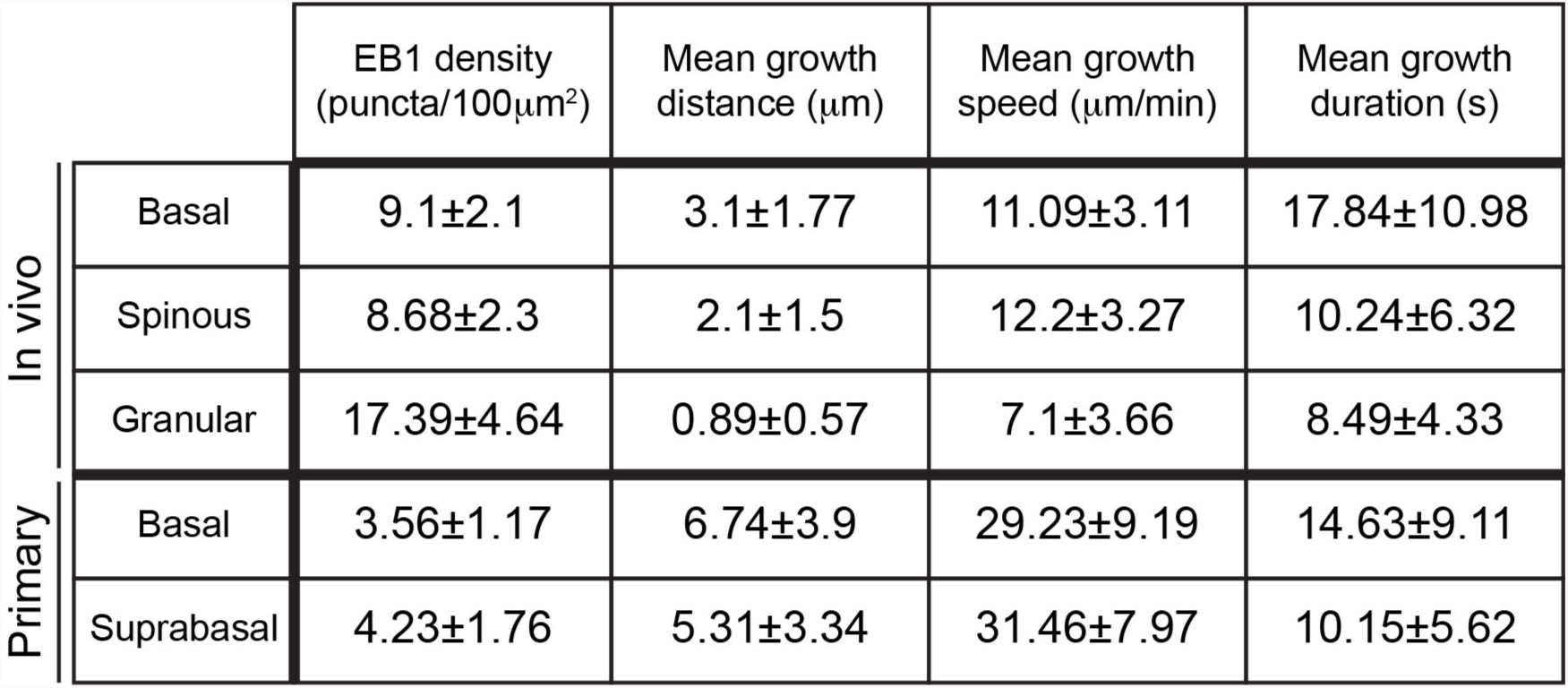
Quantifications of microtubule parameters in indicated cell types. Data are represented as mean±standard deviation. n=160 microtubules for each cell type.

**Figure 5-figure supplement 1.**
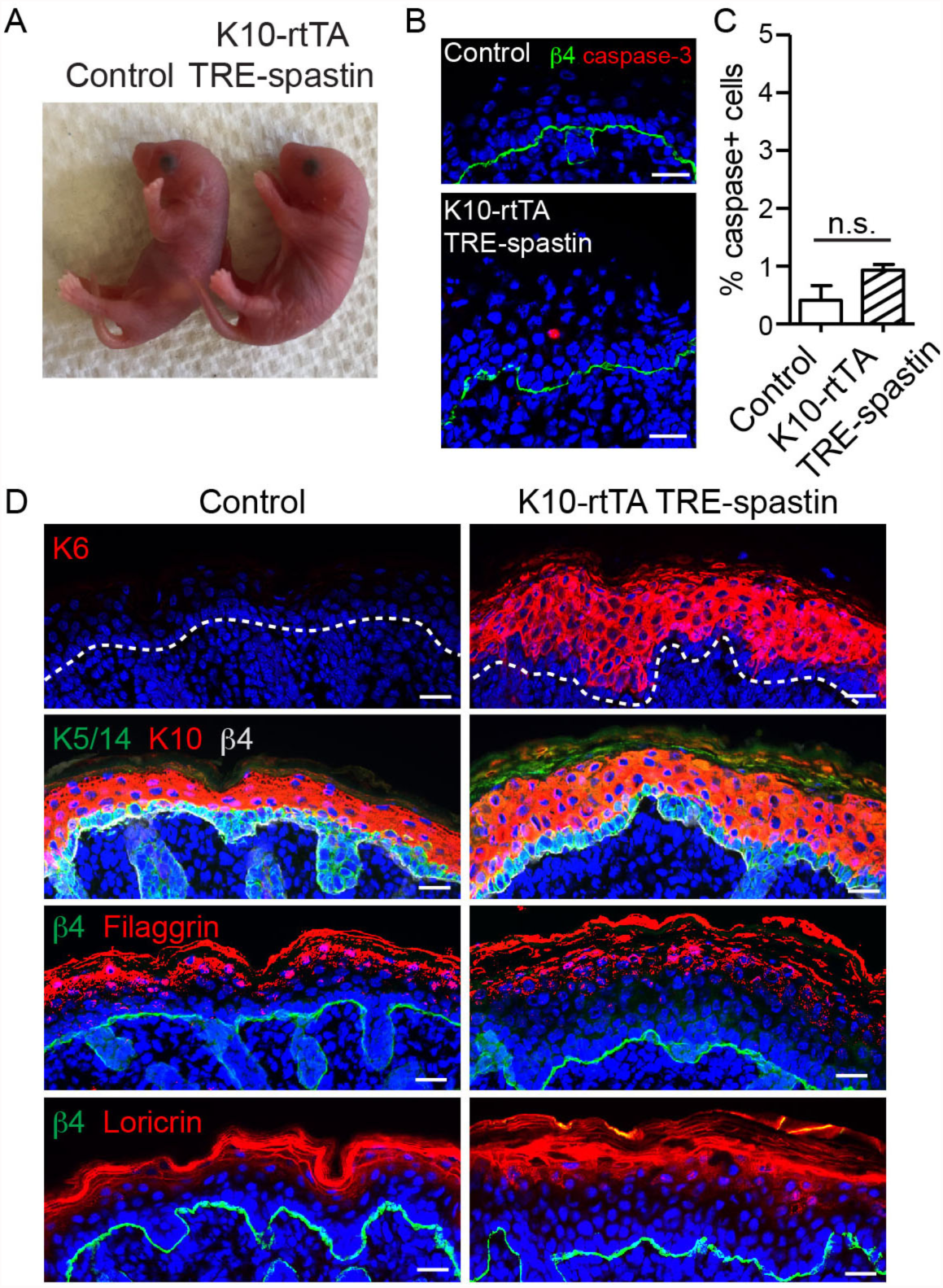
Characterization of K10-rtTA; TRE-spastin epidermis. A. P0 control and K10-rtTA; TRE-spastin mice. B. Cleaved caspase-3 staining in control and K10-rtTA; TRE-spastin epidermis. C. Quantification of cleaved-caspase-3-positive cells in control and K10-rtTA; TRE-spastin epidermis. n=4 mice for each genotype. D. Epidermal cross-sections stained for markers of stress (K6), stratification (K5/14 and K10), and terminal differentiation (loricrin and filaggrin). All scale bars are 25μm. Data are presented as mean±S.E.M. n.s.-p>0.05.

**Figure 5-figure supplement 2.**
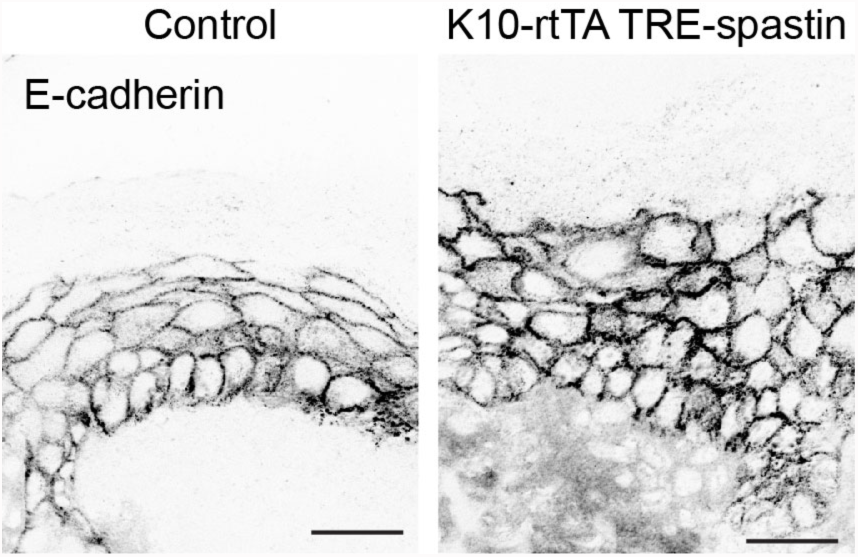
Localization of E-cadherin in control and K10-rtTA; TRE-spastin epidermis. Scale-25μm.

**Figure 5-figure supplement 3.**
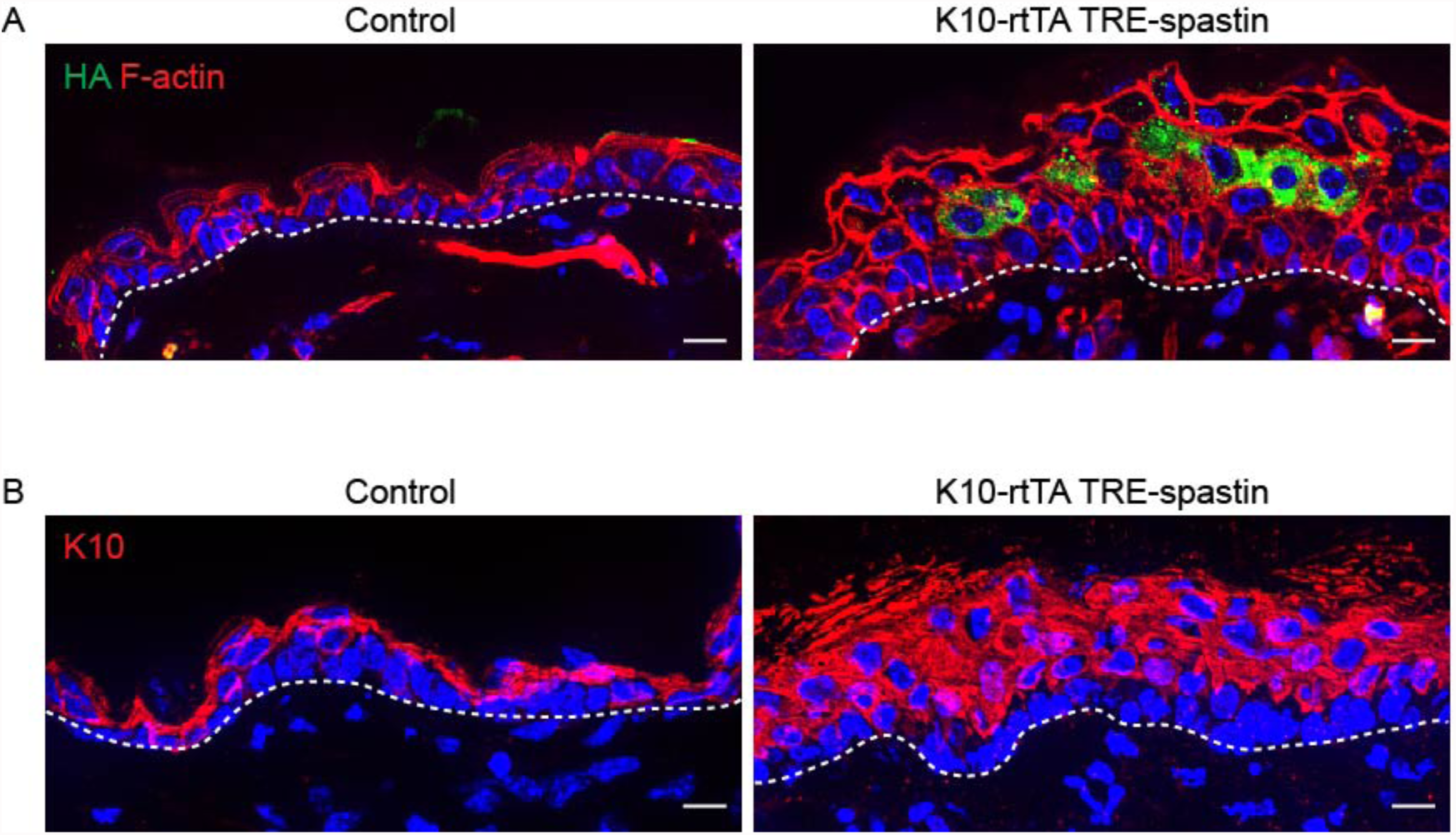
Spastin OE in suprabasal keratinocytes in adult mice perturbs epidermal homeostasis. A. Cross-section of adult (P45) epidermis in control and K10-rtTA; TRE-spastin epidermis demonstrating epidermal thickening in the mutant. B. Spastin OE in adult mice causes a thickening of the suprabasal, K10-positive layers of the interfollicular epidermis. All scale bars are 10μm.

**Figure 7-figure supplement 1.**
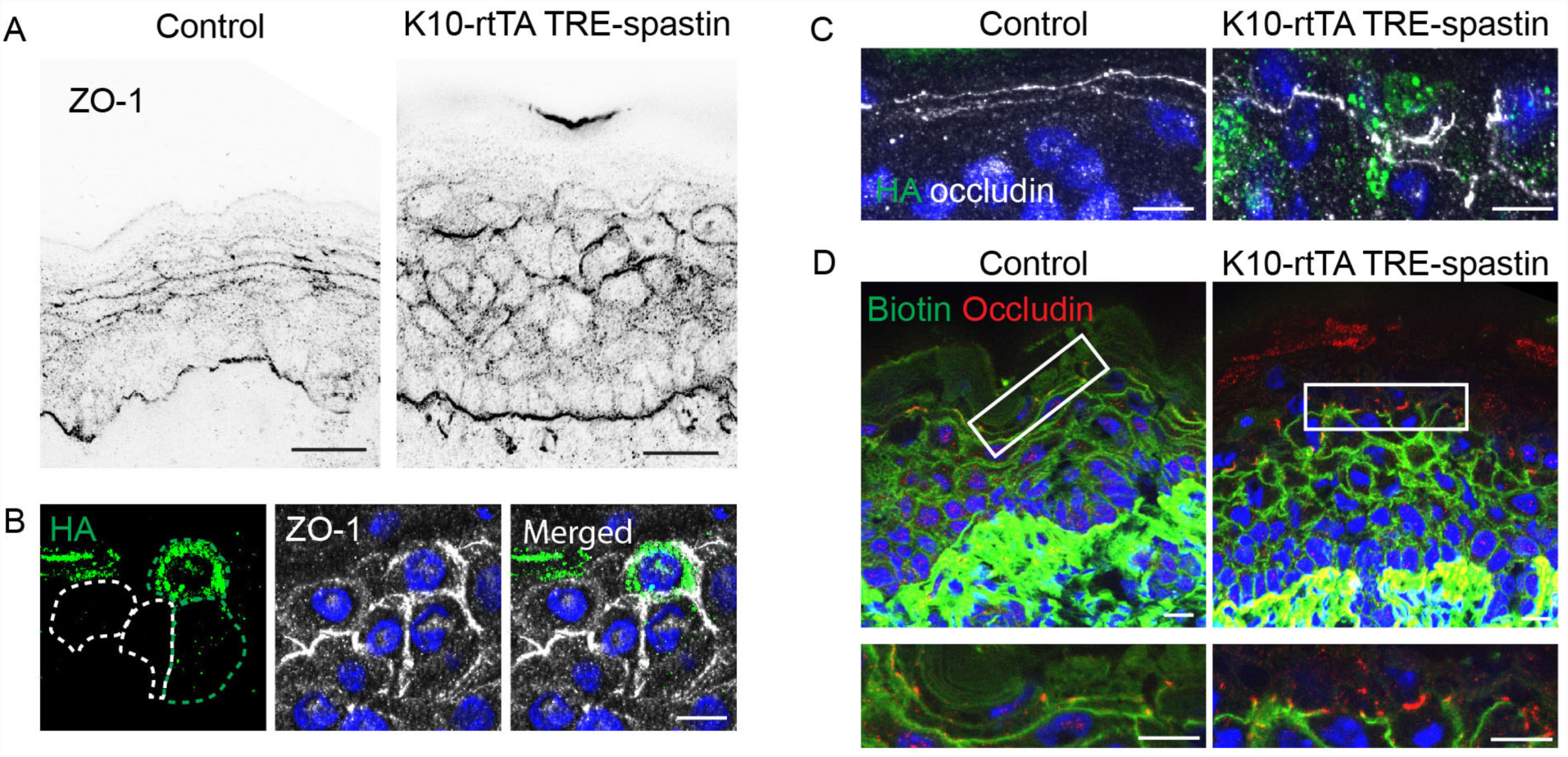
Spastin OE does not impair tight junction localization or function. A. ZO-1 localization in control and K10-rtTA; TRE-spastin epidermis. Scale-25μm. B. Region where spastin+ cells are next to spastin- cells in K10-rtTA; TRE-spastin tissue. Note that ZO-1 is still cortically localized in spastin+ cells. Scale-10μm. C. Localization of occludin at the cell cortex is maintained in K10-rtTA; TRE-spastin epidermis. Scale-10μm. D. Biotin diffusion is blocked by occludin in K10-rtTA; TRE-spastin epidermis. Scale-10μm.

## Supplemental Movie Files

Movie S1: EB1-GFP dynamics in a proliferative, basal keratinocyte in a mouse embryo.

Movie S2: EB1-GFP dynamics in a differentiated, spinous keratinocyte in a mouse embryo.

Movie S3: EB1-GFP dynamics in a differentiated, granular keratinocyte in a mouse embryo.

